# Pharmacological targeting of host chaperones protects from pertussis toxin in vitro and in vivo

**DOI:** 10.1101/2020.09.24.303321

**Authors:** Katharina Ernst, Ann-Katrin Mittler, Veronika Winkelmann, Nina Eberhardt, Anna Anastasia, Michael Sonnabend, Robin Lochbaum, Jan Wirsching, Ciaran Skerry, Nicholas H. Carbonetti, Manfred Frick, Holger Barth

## Abstract

Whooping cough is caused by *Bordetella pertussis* that releases pertussis toxin (PT) which comprises enzyme A-subunit PTS1 and binding/transport B-subunit. After receptor-mediated endocytosis, PT reaches the endoplasmic reticulum from where unfolded PTS1 is transported to the cytosol. PTS1 ADP-ribosylates G-protein α-subunits resulting in increased cAMP signaling. Here, the role of target cell chaperones Hsp90, Hsp70, cyclophilins and FK506-binding proteins for cytosolic PTS1-uptake is characterized in detail. PTS1 specifically and directly interacts with chaperones *in vitro* and in cells. Specific pharmacological chaperone inhibition protects CHO-K1, human primary airway basal cells and a fully differentiated airway epithelium from PT-intoxication by reducing cytosolic PTS1-amounts without affecting cell binding or enzyme activity. PT is internalized by human airway epithelium secretory but not ciliated cells and leads to increase of apical surface liquid. Cyclophilin-inhibitors reduced leukocytosis in infant mouse model of pertussis, indicating their promising potential for developing novel therapeutic strategies against whooping cough.

## Introduction

Whooping cough is a highly transmissible respiratory disease caused by droplet infection of the respiratory tract with *Bordetella (B*.*) pertussis*. This bacterium mediates disease through several virulence factors including adhesion factors and the enzymatically active pertussis toxin (PT) ^1^. Characteristic symptoms include severe paroxysmal coughing, which typically lasts for several weeks and can cause secondary complications like vomiting, rib fractures and pneumothorax. In severe cases, further complications such as pneumonia, encephalopathy, seizures and apnea can lead to death especially in newborns and infants ^2^. Leukocytosis, the rapid, unregulated expansion of circulating leukocytes, is a hallmark of severe infant disease ^3,4^. High levels of leukocytosis are associated with poor disease outcome and death ^3^. In 2008, whooping cough caused death in 195,000 of an estimated 16 million cases worldwide according to the world health organization (WHO) ^5^. Despite widespread vaccine coverage, disease incidence is at its highest since the 1950s ^6^.

*B. pertussis* infects humans and colonizes the ciliated epithelium of the respiratory tract. Adhesion factors like filamentous hemagglutinin (FHA) and fimbria are required for efficient infection. Moreover, *B. pertussis* produces several toxins that directly influence physiological functions. The precise role of PT for the pathogenesis of *B. pertussis* is still not completely elucidated. In mouse models, PT causes exacerbated and prolonged airway inflammation ^7^. Mutant strains of *B. pertussis* that do not express PT, do not cause severe symptoms such as leukocytosis or death in animal models ^8^. This observation clearly indicates that PT promotes disease severity and represents an attractive drug target ^7,9,10^.

PT is an AB_5_-type protein toxin, consisting of one enzyme subunit, the A protomer PTS1, and four different binding (B) subunits PTS2, PTS3, PTS4 and PTS5 occurring in the ratio 1:1:2:1, respectively ^11,12^. In *B. pertussis*, PTS1 and the B pentamer are assembled in a non-covalent manner as PT holotoxin in the periplasm prior to secretion by type VI secretion system ^11,13^. The B oligomer facilitates binding of PT to sialic acid-containing glycoproteins present on most mammalian cell types. PT does not appear to bind a single unique receptor, instead it binds in a non-saturable, non-specific manner ^14^. PT is endocytosed and follows a retrograde route of intracellular trafficking travelling through the Golgi apparatus to the endoplasmic reticulum (ER). Consequently, brefeldin A (BFA), which leads to disassembly of the Golgi apparatus, inhibits intoxication of CHO cells with PT ^15–17^. In the ER, binding of ATP to PT destabilizes interaction between PTS1 and B oligomer, which contributes to the release of PTS1 ^18,19^. ATP-mediated release occurs in the ER because it is one of the few cellular compartments besides the cytosol and lysosomes/lysosome-related organelles that contains ATP ^20^. Released PTS1 is thermally unstable and therefore in its unfolded conformation, which makes it susceptible for transport from the ER into the cytosol via the ER-associated degradation (ERAD) pathway ^21,22^. ERAD is responsible for transport of misfolded proteins from the ER into the cytosol where they become ubiquitinated and degraded by the proteasome system ^22^. PTS1 is able to utilize this transport pathway and simultaneously evade subsequent degradation since it contains no lysine residues, which are required for ubiquitination ^23^. The molecular mechanism underlying membrane transport and refolding of PTS1 and a possible role of host cell factors in this process are not sufficiently understood. In the cytosol, PTS1 covalently transfers ADP-ribose from NAD^+^ onto the α-subunit of trimeric inhibitory G proteins (Giα) ^24,25^. This ADP-ribosylation inhibits the activity of Giα as a negative regulator of membrane-bound adenylate cyclase, hence leading to increased intracellular cAMP levels and therefore disturbed signal transduction.

Consequences of cAMP increase depend on the target cells. Presence of the PT-receptor on a wide range of different cell types might explain the various effects of PT during infection with *B. pertussis*. PT inhibits recruitment of neutrophils, monocytes and lymphocytes to the respiratory tract up to one week after infection ^26,27^. PT leads to decreased amounts of pro-inflammatory chemokines and cytokines and increased bacterial burden in early stages of infection in a mouse model ^28^. The current therapeutic regimen only comprises antibiotic treatment, which is important to inhibit rapid spreading of disease via droplet infection but impacts course of disease only if administered in very early stages of infection. Moreover, antibiotic treatment does not improve symptoms including typical paroxysmal cough and in severe manifestations pneumonia, seizures, encephalopathy, apnea and death ^2^. Resurgence of pertussis disease and lack of effective treatments necessitates development of novel therapeutics for treatment of severe whooping cough. The PT-dependent nature of severe disease suggests that targeting PT should be an effective strategy for drug development.

We recently demonstrated that specific members of the cyclophilin (Cyp) family, namely CypA and Cyp40 are involved in uptake of PTS1 into the cytosol of target cells ^29^. Cyps are members of the peptidyl prolyl cis/trans isomerases (PPIases), which are protein folding helper enzymes that catalyze the rotation of peptide bonds in proteins and assist chaperones such as heat shock protein (Hsp) 90 in protein folding/refolding ^30–32^. Both CypA and Cyp40 directly bound to PTS1 *in vitro* and application of pharmacological Cyp inhibitors such as cyclosporine A (CsA) protected CHO-K1 cells from intoxication with PT. In the presence of Cyp inhibitor, less PTS1 reached the cytosol and there was less ADP-ribosylated Giα, implicating that Cyps are required for uptake of PTS1 into the cytosol ^29,33^.

Here, the role of chaperones and PPIases for the uptake of PT into target cells was investigated and Hsp90, Hsp70, as well as specific Cyps and FK05 binding proteins (FKBPs) were identified as novel interaction partners of PTS1. Application of specific pharmacological inhibitors revealed their crucial functional role for the transport of PTS1 into the cytosol of target cells, protected cells, including primary human airway cells, from intoxication with PT and reduced leukocytosis in an infant mouse model of pertussis.

## Results

### Specific pharmacological inhibition of chaperone and PPIase activities protects cells from intoxication with PT

A previously established morphology based assay of cellular intoxication was used to determine the functional role of host cell chaperones and PPIases during PT-intoxication ^29,34^. CHO-K1 cells show a characteristic clustering morphology if treated with PT (Fig 1). CHO-K1 cells were pre-incubated prior to PT-treatment with radicicol (Rad) to block Hsp90-activity, and FK506 or CsA to inhibit FKBP or Cyp activity, respectively. Presence of Rad, CsA or FK506 robustly reduced CHO-K1 cell clustering compared to cells treated with PT (Fig 1A). BFA was used as an inhibitor of PT-induced CHO-K1 cell clustering ^17,29,35^. Additionally, fewer cells were observed in wells treated with PT (Fig 1A). This effect was also inhibited by Rad, CsA and FK506. Hsp70 inhibitors VER and HA9 both inhibited PT-intoxication (Fig 1B). Taken together, the results indicate a functional role of Hsp90, Hsp70, Cyps and FKBPs for PT-intoxication.

**Figure 1.**
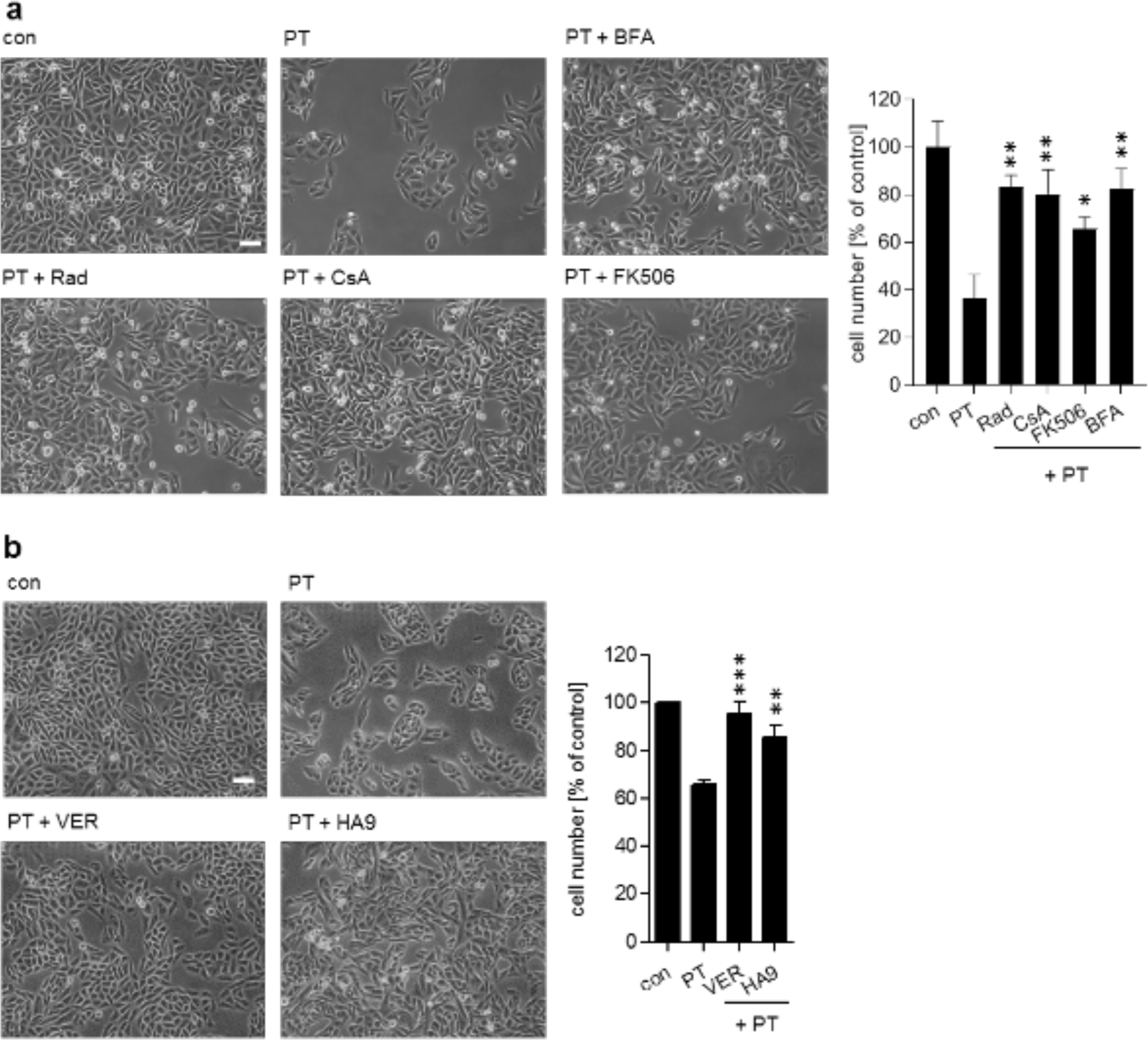
**A. Effect of BFA, Rad, FK506 and CsA on the intoxication of CHO-K1 cells with PT.** CHO-K1 cells were pre-incubated with 10 µM BFA, Rad, FK506 or CsA or left untreated for control. After 30 min 10 ng/ml PT were added. 1.5 h later the culture medium was removed, and cells were further incubated at 37 °C and 5 % CO_2_ in fresh medium that did not contain PT or any inhibitor. Pictures were taken after 18 h. A quantitative analysis of total cell number of CHO-K1 cells is shown, values are normalized on control cells (n = 3, mean ± SD). **B. Pharmacological inhibition of Hsp70 activity protects cells from PT-intoxication**. CHO-K1 cells were treated with VER (30 μM) or HA9 (20 µM) for 30 min and then intoxicated with PT (10 ng/ml) for 18 h. Pictures were taken, and cell numbers were determined as described in A. Significance was tested by student’s t-test and refers to samples treated with PT only (*p < 0.05, **p < 0.01, *** p < 0.001).

### Inhibition of chaperone/PPIase activities leads to reduced ADP-ribosylated Giα without interfering with enzyme activity of PTS1 or binding of PT to cells

After incubation of CHO-K1 cells with PT in the presence or absence of the chaperone and PPIase inhibitors, cells were lysed and incubated with fresh PTS1 in the presence of biotin-labeled NAD^+^. During this incubation, the portion of Giα, which has not been modified by PTS1 during the previous treatment of living cells with PT, was ADP-ribosylated *in vitro* ^29^. This results in biotin-labeling of Giα due to the covalent transfer of the biotin-labeled ADP-ribose moiety from NAD^+^. Biotin-labeled i.e. ADP-ribosylated Giα was then detected by Western blotting. A strong signal was obtained from control samples, whereas a weaker signal was observed in the presence of PT, showing that in PT-treated cells a substantial amount of Giα was already ADP-ribosylated and could not serve as substrate in the subsequent *in vitro* ADP-ribosylation reaction. Pre-treatment of cells with chaperone/PPIase inhibitors reduced the ADP-ribosylation of Giα compared to cells treated with PT alone (Fig 2A). Inhibition of Hsp90, Hsp70, FKBPs and Cyps prevented PT-mediated cAMP increase (Fig 2B).

**Figure 2.**
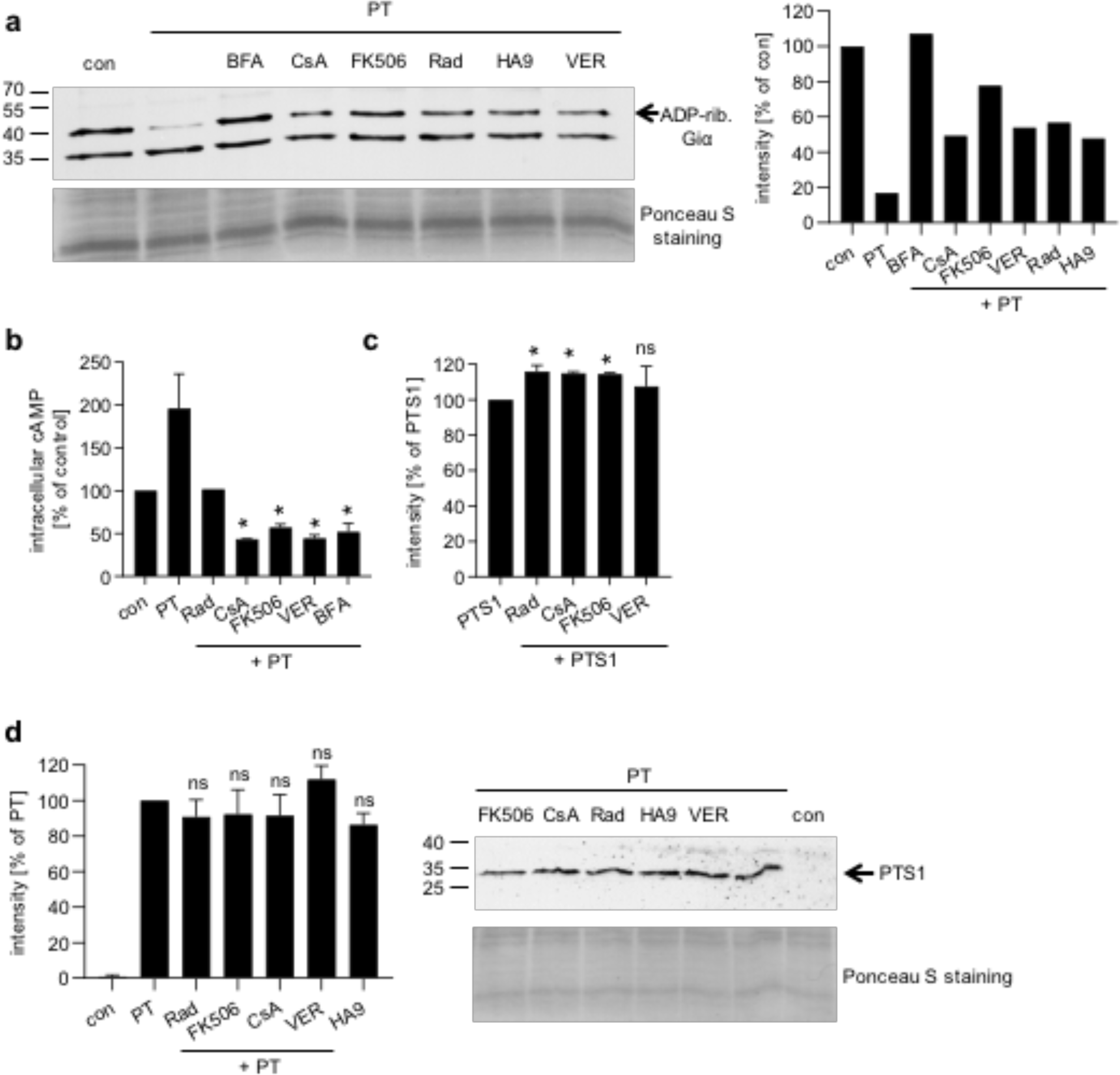
**A. Effect of Rad, CsA, FK506, VER and HA9 on the ADP-ribosylation status of Giα in PT-treated CHO-K1 cells.** Cells were pre-incubated with 20 µM of Rad, CsA or FK506, 30 µM VER, 100 µM HA9 or 20 µM BFA for control for 30 min or left untreated. Then, cells were challenged with 20 ng/ml PT for 3 h. Cells were lysed and ADP-ribosylation status of Giα was analyzed. Biotin-labeled (i.e., ADP-ribosylated) Giα was detected. Equal protein loading was confirmed with Ponceau-S-straining. Western blot signals were quantified and normalized to protein loading (Ponceau S staining). One representative result is shown (n = 3). **B. Effect of Rad, CsA, FK506 and VER on intracellular cAMP levels in PT-treated cells**. CHO-K1 cells were pre-incubation with 20 µM of Rad, CsA, FK506 or BFA or 30 µM VER for 30 min or left untreated. After 3 h of PT incubation (1 µg/ml), cells were lysed, and cAMP ELISA was performed according to the manufacturer’s manual. Values are given as percent of untreated control (n = 2, mean ± SD). Significance was tested by student’s t-test and refers to samples treated with PT only (*p < 0.05). **C. Effect of Rad, CsA, FK506 and VER on enzyme activity *in vitro***. CHO-K1 cell lysates were pre-incubated with 20 µM Rad, CsA, FK506 or 30 µM VER for 30 min or with buffer for control. After 30 min 170 ng PTS1 and 10 µM biotin-labeled NAD^+^ were added and incubated for 30 min. Samples were subjected to SDS-PAGE, blotted and ADP-ribosylated Giα was detected with Strep-POD. Western blot signals were quantified and normalized to loaded protein. Values are given as percent of samples treated with PTS1 only (n = 3, mean ± SD). Significance was tested by student’s t-test and refers to samples treated with PTS1 only (*p < 0.05, ns = not significant). **D. Influence of Rad, CsA, FK506 and VER on receptor binding of PT to CHO-K1 cells**. CHO-K1 cells were pre-incubated with 20 µM Rad, CsA, FK506 or 30 µM VER or left untreated. After 30 min cells were cooled to 4 °C for 15 min. Then 500 ng/ml PT were added for 30 min at 4 °C. After washing, cells were lysed with Laemmli buffer at 95 °C. Cell-bound PT was detected via an anti-PTS1 antibody in Western Blot. Equal protein loading was confirmed with Ponceau-S-staining. Quantification of Western blot signals and one representative Western blot is shown. Values were normalized on the amount of loaded protein and are given as percent of PT binding (second bar from left). (n = 4, mean ± SEM). Significance was tested by student’s t-test and refers to samples treated with PT only (ns = not significant).

Taken together, these results suggest that inhibition of chaperones/PPIases reduced PTS1-mediated cytosolic ADP-ribosyltransferase activity. To explain this, we have developed competing hypotheses: 1) the inhibitors reduce the enzyme activity of PTS1 or 2) the inhibitors interfere with the transport of PTS1 into the cytosol.

None of the inhibitors impaired the ADP-ribosylation of Giα by PTS1 *in vitro* (Fig 2C), suggesting that the inhibitors prevent uptake of PTS1 into the cytosol. Binding of PT was not inhibited by chaperone/PPIase inhibitors (Fig 2D). This implies that another step of toxin uptake, such as the intracellular membrane transport of PTS1 into the cytosol is the target of these inhibitors.

### In the presence of chaperone/PPIase inhibitors, less PTS1 is detectable in cells

Previously we used fluorescence microscopy to demonstrate that a monoclonal PTS1-antibody recognizes preferably PTS1 when it is detached from the B-subunit pentamer, i.e. cytosolic PTS1 ^29^. Detection of PTS1 is decreased following incubation with chaperone/PPIase inhibitors compared to cells challenged with PT only (Fig 3). This effect was also observed with BFA, which prevents transport of PT from the Golgi apparatus to the ER and the results are in line with our earlier findings that CsA inhibits uptake of PTS1 into the cytosol ^29^. Hence, the reduction of the PTS1 signal in the presence of chaperone/PPIase inhibitors strongly suggests that chaperones/PPIases are involved in translocation of PTS1 from the ER to the cytosol.

**Figure 3.**
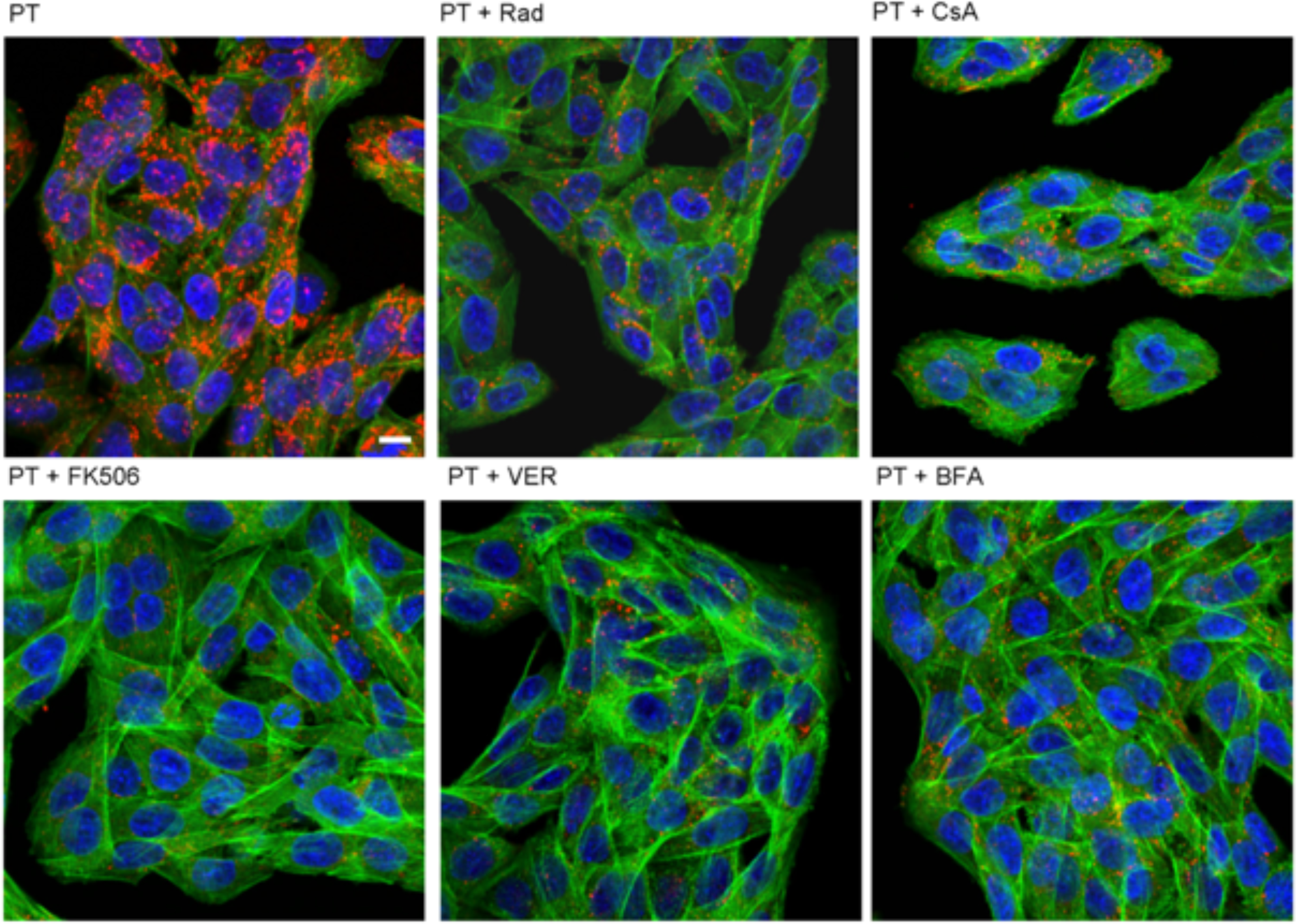
In the presence of inhibitors of Hsp90/Hsp70 and PPIases less PTS1 is detected in CHO-K1 cells. CHO-K1 cells were pre-incubated with CsA, FK506, Rad (20 µM) or VER (30 µM) for 30 min. 20 µM BFA were used as control. Then, cells were challenged with 1 µg/ml PT and 1 h later medium was exchanged. After 24 h, cells were fixed, permeabilized and blocked. Subsequently, cells were probed with an anti-PTS1 antibody, Hoechst and phalloidin-FITC for F-actin staining. Pictures were taken with a Zeiss LSM-710 confocal microscope. Red = PTS1, green = F-actin, blue = nucleus, scale bar = 10 µm.

### PTS1 directly and specifically interacts with chaperones and PPIases in vitro

To determine if PTS1 interacts directly with chaperones and PPIases, recombinant chaperones/PPIases were immobilized on a nitrocellulose membrane and overlaid with PTS1. Results in Fig 4A show specific and concentration-dependent binding of PTS1 to Hsp90, Hsp70 and Hsc70 as well as to CypA, Cyp40, FKBP51 and FKBP52. Previously, we showed that other toxin subunits like the lethal factor of *B. anthracis* or the transport component of *C. botulinum* C2 toxin do not interact with these chaperones/PPIases in the same assay indicating its specificity ^36,37^. Ponceau S-staining confirmed transfer of recombinant proteins onto the membrane. Moreover, no interaction was observed with a control protein C3bot from *C. botulinum* or the small isoform FKBP12. FKBP12 consists of only one PPIase domain. FKBP51 and FKBP52 also contain a PPIase domain and have additional domains that facilitate the interaction with potential client proteins ^32^. Here, we investigated the binding of PTS1 to FKBP-fragments that only comprise the conserved PPIase domains i.e. FKBP51 FK1 and FKBP52 FK1. Results shown in Fig 4B indicate that there still is an interaction of PTS1 with the fragments, but it is obviously weaker, compared to full-length FKBPs. This suggests that the PPIase domain, as well as the other domains of FKBPs, are involved in binding of PTS1.

**Figure 4.**
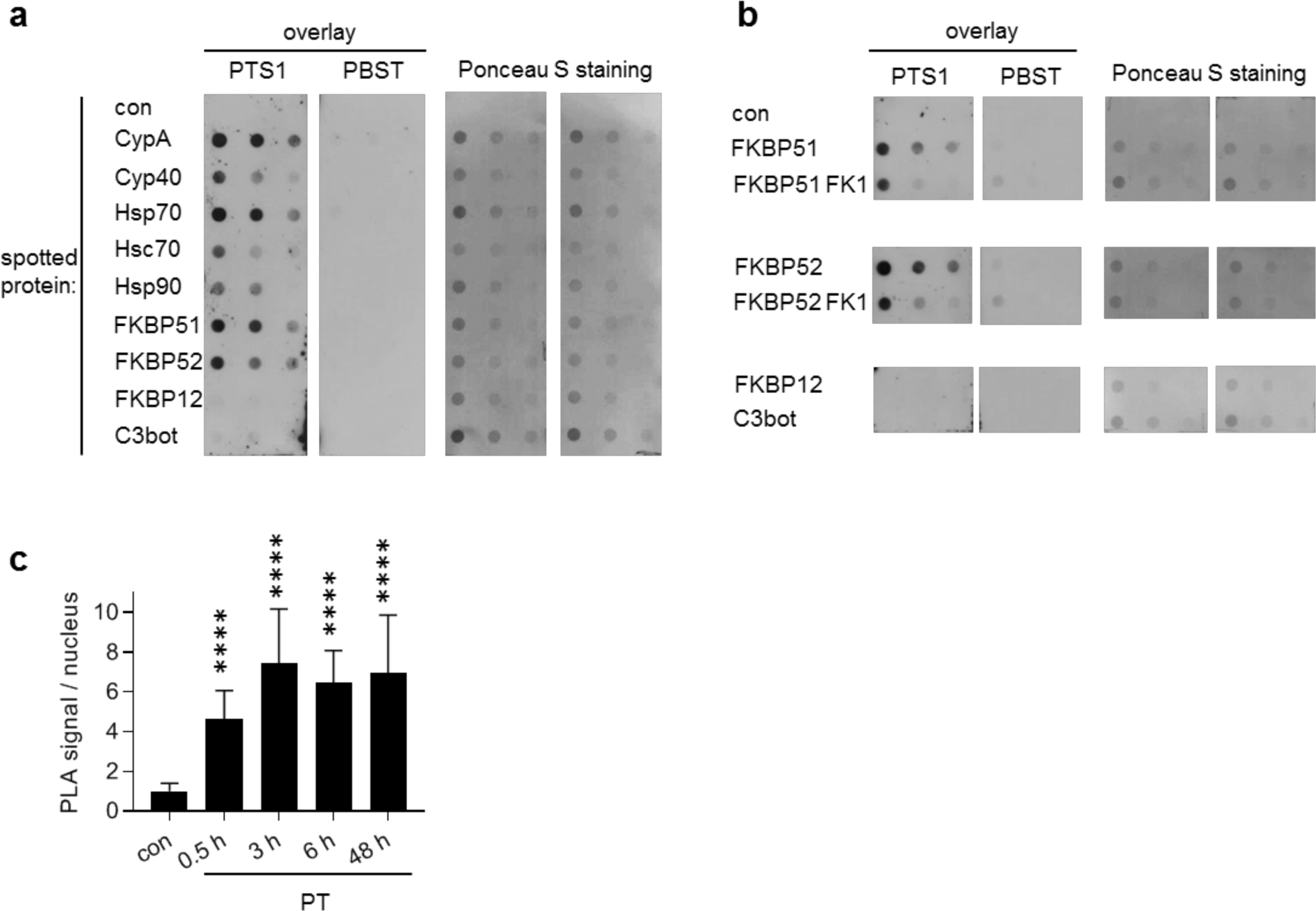
**A. PTS1 interacts with chaperones and PPIases *in vitro*.** Purified chaperones/PPIases were spotted two times onto a nitrocellulose membrane in decreasing concentrations (1 µg, 0.5 µg, 0.25 µg). PBS and C3 toxin from *C. botulinum* were used as negative controls. Membrane was cut and overlay with His-PTS1 (200 ng/ml) or PBST was performed. After extensive washing, bound PTS1 was detected with a specific antibody. PTS1- and PBST-overlayed membranes were detected on the same X-ray film and images were cropped for display purposes only. Comparable amounts of spotted protein were confirmed by Ponceau S-staining. **B. PTS1 binds to the isolated PPIase domains of FKBP51/52**. FKBP51/52 and their FK1 fragments i.e. isolated PPIase domains were spotted onto a nitrocellulose membrane. The experiment was further conducted as described in A. All signals were detected on the same X-ray film and images were cropped for display purposes only. **C. Cyp40 interacts with PTS1 in cells**. CHO-K1 cells were incubated on ice with PT (5 µg/ml) for 30 min to enable binding or left untreated for control. After washing, the cells were further incubated for 0.5 h, 3 h, 6 h and 48 h at 37 °C. Cells were fixed and fluorescence-based PLA assay was performed according to the manufacturer’s manual. PLA signals represent one protein interaction event of PTS1 and Cyp40 and were counted from fluorescence pictures (n = 10 pictures per condition) with ImageJ. Values were normalized to the mean of the control samples. Significance was tested by student’s t-test and refers to samples treated with PT only (**** p < 0.0001).

### Cyp40 interacts with PTS1 in cells for at least 48 h post intoxication

After demonstrating a direct interaction of chaperones/PPIases with PTS1 *in vitro* and establishing a functional role of these factors during uptake of PT into the cytosol, the duration of the PTS1-Cyp40 interaction was investigated in cells. The proximity ligation assay (PLA) technology encompasses an internal amplification reaction that only occurs if two antibodies against the potential binding partners come into close proximity. Every PLA signal detected by fluorescence microscopy represents one interaction event between PTS1 and Cyp40. The interaction between PTS1 and Cyp40 was detected from 30 min up to 48 h after incubation of cells with PT (Fig 4C).

### PT invades secretory but not ciliated cells in a human airway epithelium

To determine if the phenomenon of chaperone/PPIase requirement for PT uptake is recapitulated in a model of primary human bronchial airway epithelium, we first demonstrated that PT intoxicates human primary basal cells from airway epithelium in a concentration-dependent manner (supplemental Fig 1). Although clustering was not observed in these cells upon PT-treatment, impairment of cell morphology was noted (supplemental Fig 1) suggesting some cell damage induced by PT. Application of Rad or CsA reduced ADP-ribosylation of Giα in PT-treated basal cells (Fig 5A), as analyzed by sequential ADP-ribosylation. Mechanistic studies performed in CHO-K1 cells demonstrated that Rad and CsA do not influence enzyme activity or receptor binding but reduce the cytosolic amount of PTS1 in target cells, suggesting that Hsp90 and Cyps are involved in the transport of PTS1 into the cytosol of these cells.

**Figure. 5.**
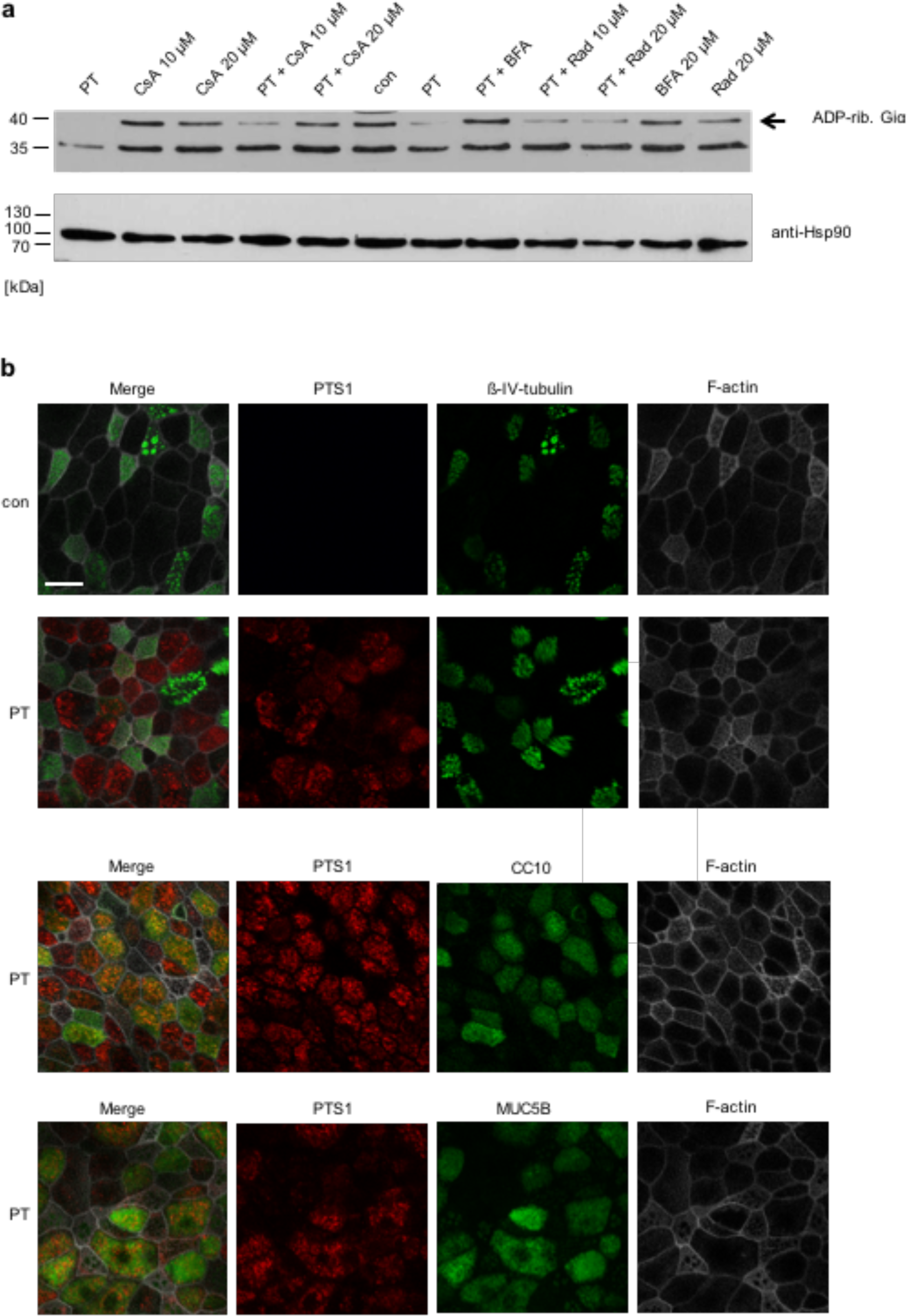
**A. Rad and CsA inhibit ADP-ribosylation of Giα in PT-treated basal cells.** Basal cells were pre-incubated with Rad, CsA or BfA (20 µM) for 45 min or left untreated for control (con). Subsequently, PT (10 ng/ml) was added for 3.5 h and the ADP-ribosylation status of Giα was determined as described before. A representative Western blot result is shown (n ≥ 2). Comparable protein loading was confirmed by Hsp90 staining. **B. PT is internalized into secretory (CC10+, MUC5B+) cells but not into ciliated (ß-IV-tubulin+) cells**. hBAECs were incubated with PT (20 µg/ml) from the apical side for at least 72 h or left untreated for control. Cells were fixed with 4 % PFA. For permeabilization and quenching of autofluorescence, cells were treated with 0.2 % saponin. F-actin was stained with phalloidin-FITC. PTS1, ß-IV-tubulin, CC10 and Muc5B were stained with specific primary and fluorescence-labeled secondary antibodies, respectively. Pictures were taken with an inverted confocal microscope. Scale bar = 20 µm.

Basal cells that represent one cell type of human bronchial airway epithelial cells (hBAECs) can be differentiated to a functional airway epithelium at air-liquid interface conditions, containing ciliated and secretory cells. By analyzing the effect of PT on the differentiated cell layer by fluorescence microscopy, it became evident that the toxin was selectively internalized in secretory (CC10 or MUC5B positive) cells ^38^, but not into ciliated cells (ß-IV-tubulin positive) (Fig 5B). Moreover, CsA but not Rad or FK506 reduced the amount of PTS1 detected by fluorescence microscopy in this functional airway epithelium (Fig 6). Rad, CsA or FK506 alone had no adverse effects on morphology of the airway epithelium or tight junctions (supplemental Fig 2A). However, VER and BFA revealed strong adverse effects on the airway epithelium showing a reduced cell number for VER-treated cells and nearly no cells were detected after BFA treatment (supplemental Fig 2B).

**Figure. 6.**
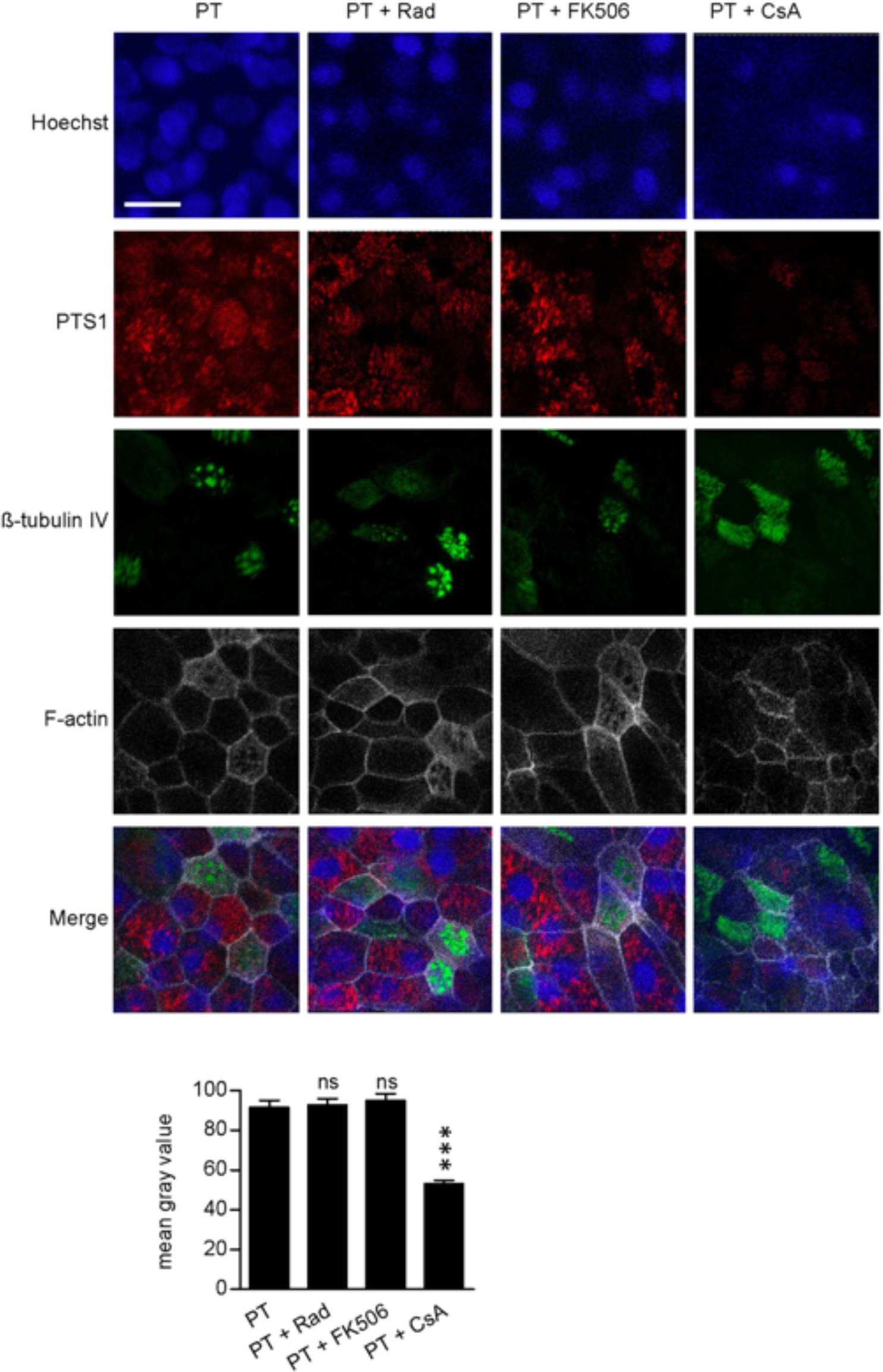
**Effect of Rad, FK506 and CsA on uptake of PTS1 into hBAECs.** hBAECs were pre-treated with respective inhibitors (20 µM) and then incubated with PT (20 µg/ml) from the apical side for 72 h. Cells were fixed with 4 % PFA. For permeabilization and quenching of autofluorescence, cells were treated with 0.2 % saponin. F-actin was stained with phalloidin-FITC. PTS1 and ß-IV-tubulin were stained with specific primary and respective fluorescence-labeled secondary antibodies. Pictures were taken with a Cell Observer inverse microscope (Zeiss, Germany). Mean Gray Value Intensity was measured and plotted for PTS1 (after background subtraction) in PT and PT + inhibitor treated cultures. Values are given as mean ± SEM (n= 60 cells/condition). Significance was tested by Kruskal-Wallis test and refers to samples treated with only PT (***p < 0.001, ns = not significant). Scale bar = 20 µm.

Next, effects of PT on functional aspects of the human bronchial airway epithelium were investigated. PT showed no significant effect on trans-epithelial electrical resistance (TEER) of the human bronchial airway epithelium (Fig 7A). However, PT significantly increased the apical surface liquid (ASL) of the functional human airway epithelium compared to untreated controls (Fig 7B). This effect was not impaired by the chaperone inhibitors suggesting that other mechanisms might play a role that do not require PTS1 activity in the cytosol such as binding of the B-pentamer to the cell surface. Chaperone/PPIase inhibitors alone led to no significant effects on ASL or TEER values of treated samples compared to control (Fig 7, right panels).

**Figure. 7.**
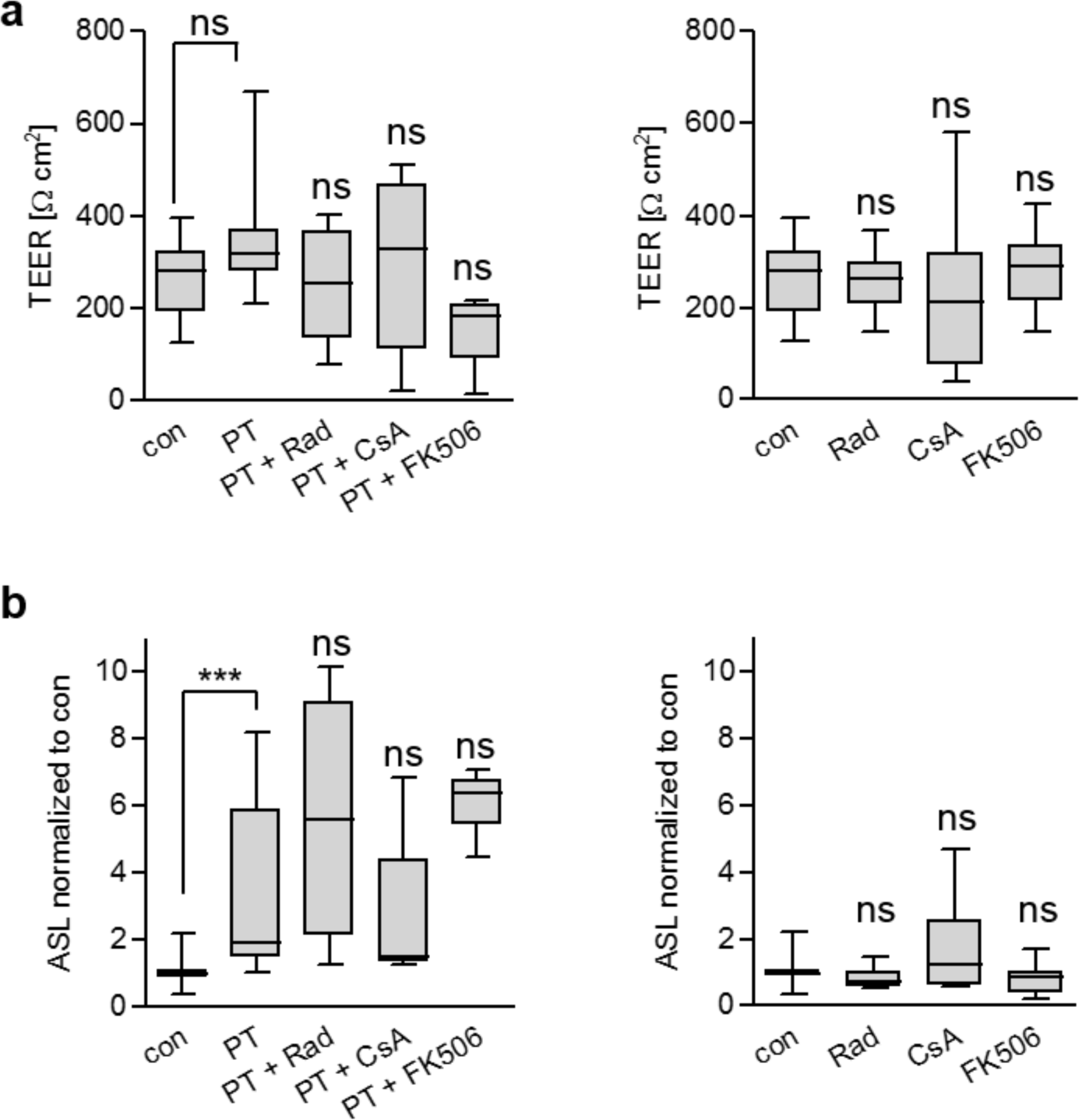
PT does not impair barrier function but increased apical surface liquid. **A**. hBAECs were pre-incubated with Rad, CsA or FK506 (20 µM) for 30 min or left untreated. Then 20 µg/ml PT was added for 72 h or cells were left untreated. Transepithelial electrical resistance (TEER) was measured using CellZScope from NanoAnalytics. **B**. hBAECs were treated as described in A. After 72 h the apical surface liquid was determined. Values were obtained from at least two independent experiments (n ≥ 4). Significance was tested by Kruskal-Wallis test and refers to samples treated with only PT or as indicated (***p < 0.001, ns = not significant).

### Cyp inhibitors reduce leukocytosis in an infant model of pertussis disease

Finally, effects of CsA and its non-immunosuppressive derivative NIM811 were investigated in an infant mouse model of pertussis. This model has recently been established and recapitulates several hallmarks of severe disease observed in humans such as leukocytosis and death, which are both PT-dependent ^8^. Here, 7-day old mice were infected with a PT-producing wild type strain of *B. pertussis* via aerosol and then treated intranasally with either CsA or NIM811 or vehicle. Treatment with CsA or NIM811 had no effect on the colony forming units detected from lung homogenates (Fig 8). However, PT-induced leukocytosis was significantly reduced by CsA or NIM811 treatment in infant mice implicating a protective effect upon Cyp inhibition for the first time *in vivo*.

**Figure. 8.**
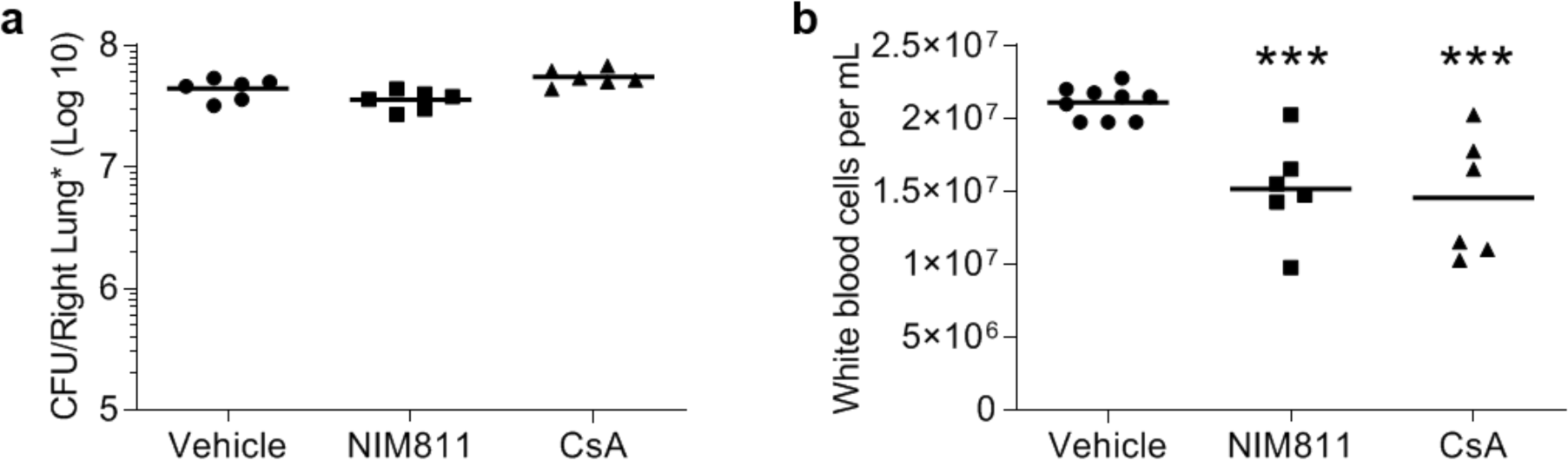
Cyclophilin inhibition reduces *B. pertussis* induced leukocytosis. Following aerosol infection 7-day old C57BL/6 pups were treated daily with vehicle, CsA (25 mg/kg) or NIM811 (25 mg/kg). No differences were determined in bacterial burden **(A)** however significant reduction in leukocytosis was noted in drug treated animals versus control **(B)**. Each data point represents one animal. Significance was determined by students t-test ***=p<0.001. Bars represent mean values.

## Discussion

Pertussis disease is a reemerging public health crisis for which there is currently no effective treatment. The current standard of care, treatment with macrolide antibiotics is only associated with improved symptoms when administered during the catarrhal stage of disease ^39^. This is challenging as it is before the onset of the characteristic “whooping” cough. *B. pertussis* mediates disease through its ADP-ribosylating toxin (ADP-RT) PT. The ability of ADP-RTs to translocate their enzyme subunits to target cell cytosols has been the subject of much research (for review see ^40,41^). We have shown other membrane translocating ADP-RTs, such as *Corynebacterium diphtheriae* diphtheria toxin, *C. botulinum* C2 toxin, *C. perfringens* iota toxin, and *C. difficile* CDT, require the chaperone activity of heat shock proteins Hsp90 and Hsp70, Cyps as well as PPIases from the FKBP family to transfer their enzymatic subunits from vesicular compartments to the cytosol of target cells ^37,42–50^. Identifying commonalities in these toxins mechanisms of action may allow for the development of treatments which function for across diverse bacterial species.

Our previous work identified Cyp inhibitors as potent inhibitors of PT intoxication of mammalian cells ^29^. Previously, we determined that Cyp isoforms CypA and Cyp40 interact with PTS1 *in vitro* and are required for translocation of PTS1 into the cytosol. This study revealed host Hsp90, Hsp70, Cyps and FKBPs are required for translocation of PTS1 from the ER to the cytosol. A direct and specific interaction of PTS1 with Hsp90, Hsp70, CypA, Cyp40, FKBP51 and FKBP52 was detected *in vitro* and interaction with FKBPs was partly mediated by the PPIase domain of these proteins. Human bronchial derived airway epithelial cells differentiated under air-liquid interface conditions allowed us to identify secretory cells as targets for PTS1 intoxication. Further, pharmacological inhibitors of Cyps protected against leukocytosis in an *in vivo* murine model of infant pertussis disease demonstrating the clinical potential of these molecules.

ADP-ribosylation of GTP-binding regulatory protein Giα by PT ^12^ induces distinct morphological changes in CHO-K1 cells in a sensitive and reproducible manner ^34^. This phenomenon is the basis of routine assays to determine monoclonal antibody function and to measure antitoxin responses ^51,52^. Here, we utilized this CHO-K1 assay and pharmacological inhibitors of heat shock proteins, Cyps and FKBPs to show PT interactions with Hsp90, Hsp70, FKBP and Cyps are required for PT-intoxication of CHO-K1 cells. Further, we determined inhibition of these host molecules prevents ADP-ribosylation of Giα protein and intracellular cAMP accumulation of CHO-K1 by preventing translocation of PT from the endoplasmic reticulum to the cytosol, not by inhibiting PT enzymatic activity. Immobilization of the studied host chaperones and folding helper proteins on a nitrocellulose membrane and incubation with PTS1 identified Hsp90, Hsp70 and Hsc70, CypA, Cyp40, FKBP51 and FKBP52 as binding partners of PTS1 in a specific and concentration-dependent manner. Cyp40, FKBP51 and FKBP52 contain three TPR domains through which they bind to Hsp90 ^32,53^ forming a Hsp90-multichaperone complex ^30,31^. The Hsp90-multichaperone complex facilitates the stabilization, folding and activation of more than 200 mammalian proteins ^54–56^.

Recently, it has been shown that cholera toxin requires Hsc/Hsp70 but not Cyps in addition to Hsp90 for its cellular uptake ^57^. However, interaction of Hsc/Hsp70 and Hsp90 with CTA1, the enzyme domain of cholera toxin, occurs independently since both chaperones can bind simultaneously to CTA1, and binding occurs in the absence of the adapter protein Hop. This suggests an alternative interaction mechanism between cholera toxin and Hsp90/Hsc/Hsp70 that differs from the described Hsp90-multichaperone machinery.

We showed that interaction between Cyp40 and PTS1 was detected after 48 h after PT incubation. This suggests that Cyp40 not only facilitates the translocation from the ER to the cytosol but is required to stabilize PTS1 in an active conformation. A comparable mechanism is described for the cholera toxin, which also harbors ADP-ribosyltransferase activity and belongs to the group of long trip toxins. Hsp90 facilitates translocation of CTA1 to the cytosol, refolds CTA1 into an active conformation in an ATP-dependent manner and continues to bind to refolded CTA1 ^58,59^. The finding that CTA1 and PTS1 require the assistance of chaperones even after translocation and refolding might be due to the fact that both enzyme subunits are thermally unstable. This means that unfolding of CTA1 and PTS1 occurs at 37 °C after dissociation from the B-subunit. Therefore, releasing of toxin subunits from chaperones after translocation/refolding could again result in unfolding. Recently, two Hsp90 binding motifs in CTA1, which both are necessary for efficient translocation of CTA1 into the cytosol were identified ^60^. One of these binding motifs was also identified in other ER-translocating ADP-ribosylating toxins including PT and Hsp90 requirement for PT uptake into the cytosol was demonstrated confirming and supporting the findings in the present study. Interestingly, the identified Hsp90-binding motif was absent in ADP-ribosylating toxins that translocate from endosomes suggesting an interaction mechanism distinct to some extent between short trip and long trip ADP-ribosylating toxins.

*B. pertussis* colonizes its host through attachment to the ciliated cells of the airways. It is known that PT can bind to various sialic acid-containing glycoproteins and various receptors have been found ^11,13^. However, so far it is not clear which cell types are the target for PT in humans. Human bronchial airway epithelial basal cells differentiated in air-liquid interface conditions recapitulate the characteristics of the airway epithelial barrier. These epithelial cells differentiate into pseudostratified, polarized cells including ciliated and goblet (secretory) cells. From this, we determined that CC10 or MUC5B positive secretory cells, and not ß-IV-tubulin positive ciliated cells were the target of PTS1 intoxication. It was determined that pharmacological inhibition of Cyps, but not Hsp90 or FKBP reduced detectable cytosolic PTS1 in these cells. Moreover, PT had no effect on the trans-epithelial resistance but increases the ASL of the differentiated human airway epithelium. These data suggest that PT does not impact gross epithelial barrier integrity but rather affects vectorial ion or fluid transport. The ASL shields the mucosal surface of lung epithelia and forms the first line of defense against airborne noxae and pathogens. A major task of lung epithelial cells is therefore to tightly regulate the ASL volume to maintain appropriate lung function and to ensure mucocilliary clearance. ASL volume homeostasis is regulated by balancing water secretion and resorption. Secretion is driven by apical chloride channels (e.g. CFTR), whereas the epithelial Na+ channel (ENaC) mediates resorption. Hence, our observation that PT disturbs ASL homeostasis might reflect an impact on vectorial ion transport, either activating apical chloride channels or inhibiting ENaC. It has been demonstrated that PT intoxication results in increased intracellular cAMP levels, and CFTR is activated by intracellular cAMP ^61^. Yet, future studies, including Ussing chamber measurements, will be required to decipher the impact of PT intoxication on transepithelial ion/water transport in more detail.

Results from another previous study in mice showed an upregulation of the epithelial anion exchanger pendrin in the lung after infection with *B. pertussis* ^62^. Pendrin knockout mice showed reduced lung inflammatory pathology but even higher bacterial loads during infection suggesting a role of pendrin in the pathology. Pendrin exports bicarbonate to the ASL and thereby increases pH, which possibly contributes to promoting the inflammatory pathology. These results indicate that manipulation of ASL parameters like volume or pH might be mediated by PT and play a role for the pathology and course of disease.

The mouse model of pertussis disease recapitulates the age-dependent nature of severe pertussis illness. Infant mice succumb to lethal infection at doses tolerated by adult mice. Leukocytosis, the rapid increase in vascular leukocytes, correlates with disease severity and lethality and is PT-mediated ^10,63^. Infant mice, like humans, show robust increases in circulating leukocytes following infection. Here, we showed that inhibition of Cyp activity by CsA, a licensed drug with well-known pharmacokinetics and safety profiles ^64,65^, and its non-immunosuppressive derivative NIM811 ^66^ led to significantly decreased leukocytosis upon infection with PT-producing *B. pertussis* strains. To our knowledge, this is the first time that Cyps were pharmacologically targeted to successfully prevent cytotoxic effects of a bacterial AB-type toxin in an *in vivo* model. Together with our detailed mechanistic studies in CHO-K1 cells and *in vitro* differentiated human airway epithelial model, these results provide a promising basis for the development of novel therapeutic strategies to prevent severe symptoms like leukocytosis caused by PT during *B. pertussis* infection.

## Methods

### Protein expression and purification

Recombinant proteins were expressed and purified as described before: Hsp70, Hsc70 ^42^, Hsp90 ^48^, FKBP12, FKBP51, FKBP52 ^49^, CypA, Cyp40 ^37^, FK1 domains of FKBP51 and FKBP52 ^50^ and C3bot ^67^.

### Protein-protein interaction studies via dot blot system

Purified recombinant chaperones, PPIases and PPIase fragments were vacuum aspirated onto a nitrocellulose membrane as serial dilution starting with 1 µg/ml using the dot blot system (Bio-Rad, Feldkirchen, Germany). Recombinant C3 toxin from *C. botulinum* (C3bot) was chosen as a random protein to exclude unspecific protein-protein interactions. Successful transfer was confirmed by Ponceau S staining. Membrane was blocked with 5 % skim milk powder in PBS-Tween (PBST) and then cut and probed with PTS1 (Aviva Systems, San Diego, California, USA) or for control with PBST. After extensive washing, both membranes were incubated with anti-PTS1 antibody (Santa Cruz, Heidelberg, Germany) to detect bound PTS1 using HRP-coupled secondary antibody (Santa Cruz) in combination with ECL (enhanced chemiluminescence, Millipore, Merck, Darmstadt, Germany) system. Signals for both membranes were detected on the same X-ray film. Pictures were cropped for display purposes only.

### Cell culture and intoxication experiments

Cell culture materials were obtained from Gibco unless indicated otherwise. Chinese hamster ovary cells strain K1 (CHO-K1, from DSMZ, Braunschweig, Germany) were cultivated in DMEM and HAM’s F12 containing 5 % heat-inactivated fetal calf serum (Invitrogen, Thermo Fisher Scientific, Waltham, Massachusetts, USA), 1 mM sodium-pyruvate and Penicillin-Streptomycin (PenStrep) (1:100) (Thermo Fisher Scientific). Cells were grown at 37 °C and 5 % CO_2_ as described before ^68^.Cells were trypsinized and reseeded every two to three days for at most 15 to 20 times. For intoxication experiments ^43,45,68^, cells were seeded in culture dishes and the specific pharmacological inhibitors radicicol (inhibitor of ATP-binding site of Hsp90, Sigma-Aldrich, Merck, Darmstadt, Germany), cyclosporine A (inhibitor of Cyp activity, Sigma-Aldrich, Merck), FK506 (inhibitor of FKBP activity, Sigma-Aldrich, Merck), VER (inhibitor of ATP-binding site of Hsp70, Hsc70 and Grp78, Tocris Bioscience, Wiesbaden-Nordenstadt, Germany), HA9 (inhibitor of the substrate binding domain of Hsp70) and brefeldin A (disrupts Golgi apparatus, Sigma-Aldrich, Merck) were added for 30 min. Then, PT (Sigma-Aldrich, Merck) was added either for 1 h with subsequent removal and further incubation for 18 h at 37 °C or PT was added for 18 h. Morphology of cells was documented by using a Zeiss Axiovert 40CFL microscope with a Jenoptik ProgRes C10 CCD camera. At least 3 images per treatment were taken and analyzed and the experiment was repeated independently at least 3 times. To quantify toxin-induced effects on CHO-K1 cells total number of cells per picture was manually counted using ImageJ (National Institutes of Health, Bethesda). Untreated control cells were set as 100 % and values of other samples were calculated accordingly. Materials for cell culture experiments were obtained from TPP Techno Plastic Products (Trasadingen, Schweiz).

### Primary human bronchial airway epithelial cell culture

Frozen primary human bronchial airway epithelial cells (hBAECs, Epithelix, Geneva, Switzerland) were obtained at passage 1. Of all cell types that are obtained from a human airway epithelium, only the basal cells survive. Basal cells were proliferated in growth medium (Promocell, Heidelberg, Germany) supplemented with Penicillin-Streptomycin (PenStrep, Thermo Fisher Scientific) until reaching 80% confluence in a T75 flask. Growth medium was replaced every two days. Cells were detached using DetachKIT (Promocell). 3.5 × 10^4^ cells were seeded onto a transwell filter (Corning Costar 3470; Corning, Wiesbaden, Germany) pre-coated with collagen solution (StemCell Technologies, Cologne, Germany). Cells were maintained with 200 µL growth medium in the apical and 600 µl of growth medium in the basolateral compartment. To obtain a fully differentiated epithelium containing secretory and ciliated cells, apical medium was removed after 72 h and basolateral medium was switched to differentiation medium. Differentiation medium was a 50:50 mixture of DMEM-H without glutamine and pyruvate and LHC basal (both from Thermo Fisher Scientific) supplemented with supplement pack (Promocell), PenStrep and different trace elements as described in detail before ^69^. Cells were grown until day 25 day with the basolateral medium being changed every three days. From day 14 onwards, cells were washed on the apical side every three days for 30 minutes with Dulbecco’s phosphate buffered solution (DPBS, Biochrom, Berlin, Germany) to remove accumulated mucus. hBAECs cultures from 2-3 different donors were used for each experiment.

### Sequential ADP-ribosylation of Giα in lysates from toxin-treated cells

CHO-K1 cells were pre-incubated with respective inhibitors and then intoxicated with PT for given incubation periods. Cells were lysed in ADP-ribosylation buffer (0.1 mM Tris-HCL (pH 7.6), 20 mM DTT and 0.1 µM ATP) plus protease inhibitor complete (Roche, Basel, Switzerland) as described earlier ^17,29^, followed by incubation with 170 ng PTS1 and 10 µM biotin-labeled NAD^+^ (Trevigen, Gaithersburg, Maryland, USA) for 1 h at room temperature for *in vitro* ADP-ribosylation of Giα, which had not yet been ADP-ribosylated by PT during the previous incubation. Samples were subjected to SDS-PAGE, blotted and ADP-ribosylated, i.e. biotin-labeled, Giα detected with streptavidin-peroxidase (Strep-POD, Sigma-Aldrich, Merck) using the ECL system. Equal amounts of protein were confirmed by Ponceau-S-staining. Densitometric quantification of Western blot signals was measured using Adobe Photoshop CS6 and values were normalized on the amount of loaded protein.

### cAMP measurement

Analysis of intracellular cAMP levels was performed by ELISA (acetylated format) according to the manufacturer’s manual (Enzo Life Sciences, Lörrach, Germany).

### In vitro enzyme activity of PTS1

Post nuclear supernatant (PNS) from CHO-K1 was gained from confluently cultivated CHO-K1 cells which were lysed in ADP-ribosylation buffer (0.1 mM Tris-HCL (pH 7.6), 20 mM DTT and 0.1 µM ATP) as described earlier ^17,29^, and centrifuged at 3,000 g at 4 °C for 5 min. CHO-K1 PNS (40 µg of protein) was then incubated for 30 min with 10 µM Rad, CsA, FK506 or 30 µM VER. As a control PNS was incubated with buffer only. After 30 min 170 ng PTS1 were added together with 10 µM biotin-labeled NAD^+^ and incubated for 1 h at room temperature. Samples were subjected to SDS-PAGE and blotted onto a nitrocellulose membrane. ADP-ribosylated, i.e. biotin-labeled, Giα was detected with Strep-POD. Equal amounts of protein were confirmed by Ponceau-S-staining.

### Toxin binding assay

CHO-K1 cells were pre-incubated with 10 µM Rad, CsA, FK506 or 30 µM VER. The assay was performed as previously described ^43,68^.Cells were cooled down to 4 °C for 15 min to prevent endocytosis. 500 ng/ml PT were added, and cells were incubated for 30 min at 4 °C. Subsequently, culture medium was removed, and cells were washed with PBS to remove unbound PT. Then cells were scraped with boiling SDS-sample buffer and heated for 10 min at 95 °C. SDS-PAGE and Western Blotting was performed. To detect cell-bound PT an antibody against the S1 subunit of PT and a peroxidase-coupled secondary antibody were used.

### Immunofluorescence

Fluorescence microscopy was performed as described earlier ^43,68^. Cells were fixed after intoxication experiments with 4 % paraformaldehyde (PFA), permeabilized with Triton-× 100 (0.4 % in PBS), treated with glycine (100 nM in PBS) and blocked with 5 % skim milk powder or 10 % normal goat serum (Jackson ImmunoResearch, West Grove, Pennsylvania, USA) and 1 % BSA in PBST for 1 h at 37 °C. Samples were incubated with anti-PTS1 antibody (Santa Cruz, 1:50 diluted in 10 % NGS and 1 % BSA in PBST) for 1 h at 37 °C. After washing, fluorescence-labeled secondary antibody anti-mouse 568 (goat) (Invitrogen) was added (1:750 diluted in 10 % NGS and 1 % BSA in PBST) for 1 h at 37 °C. Nuclei were stained with Hoechst and F-actin with phalloidin-FITC (Sigma-Aldrich, Merck). Between the individual working steps, cells were washed with PBS. Fluorescence imaging was performed with the iMIC Digital Microscope (FEI Munich, Germany) using the Live Acquisition 2.6 software (FEI Munich) and processed with ImageJ 1.4.3.47 software (National Institutes of Health, Bethesda).

hBAECs grown on transwell filters were fixed for 15 min in 4% paraformaldehyde in DPBS. Cells were then permeabilized and blocked for 5 min in DPBS containing 0.2% saponin and 10% FBS (Thermo Fisher Scientific). Cells were stained with primary (1:25-50) and secondary (1:400) antibodies in DPBS, 0.2% saponin and 10% FBS. The α-β-IV rabbit monoclonal antibody (ab179509) and ZO-1 (ab99462) were purchased from Abcam (Cambridge, United Kingdom), α-club cell secretory protein rat monoclonal antibody (MAB4218) was from R&D Systems (Minneapolis, Minnesota, USA) and the α-Muc5B rabbit polyclonal antibody (HPA008246) was from Sigma-Aldrich. Fluorescently labeled secondary antibodies were obtained from Molecular Probes (Thermo Fisher Scientific). Alexa Fluor 647-Phalloidin (8940) was obtained from NEB. Images were taken on an inverted confocal microscope (Leica TCS SP5, Leica, Wetzlar, Germany) using a 63x lens (Leica HCX PL APO lambda blue 63.0×1.40 OIL UV). Images for the blue (DAPI), green (AlexaFluor 488), red (AlexaFluor 568) and far-red (AlexaFluor 647) channels were taken in sequential mode using appropriate excitation and emission settings.

### Protein interaction analysis in cultured cells by Duolink using proximity ligation assay (PLA)

CHO-K1 cells were incubated with PT for indicated periods of time. Cells were fixed with 4% PFA, permeabilized and blocked with 10 % NGS and 1 % BSA in PBST. Subsequently, cells were incubated with mouse anti-PTS1 (Santa Cruz) and rabbit anti-Cyp40 (Thermo Fisher Scientific) antibodies for 1 h at 37 °C. PLA assay was performed according to the manufacturer’s protocol (Duolink using PLA technology, Sigma-Aldrich, Merck) and as described before ^43,44^.

### Measurement of the transepithelial electrical resistance (TEER)

TEER was analyzed by impedance spectrometry using the cellZscope (NanoAnalytics, Münster, Germany). For measurements, the basal electrode was overlaid by 500 µl equilibrated DMEM-H medium filters inserted and 250 µl ediuDMEM-H medium was added to the apical side of the filter. Measurements were performed immediately after positioning of apical electrodes. Data were acquired and analyzed using the software provided with the instrument (NanoAnalytics).

### Measurement of apical surface liquid (ASL) volume

ASL measurements were performed using the D_2_O dilution method as described previously ^70^. Filters with confluent cell layers were placed in 500 µl of differentiation medium containing respective inhibitors and pre-incubated for 30 min at 37°C. 25 µl of isotonic NaCl, containing respective inhibitors, was added to the apical compartment. Silicon sealed control filters loaded with 25 µl isotonic NaCl solution were randomly distributed throughout the plate to estimate volume changes caused by evaporation. After 72 h apical fluid volumes were collected with 25 μl D_2_O containing 0.9% (w/v) NaCl for analysis. Water concentrations were measured by attenuated total reflexion mid-infrared spectroscopy on A Vertex 70 FT-IR spectrometer, equipped with a BioATRCell-II unit and a liquid nitrogen cooled MCT detector (Bruker Optics, Fällanden, Switzerland), to calculate ASL volumes. Data acquisition and processing was performed using OPUS 6.5 (Bruker Optics). Before each measurement a background spectrum of the empty ATR-unit was collected and a calibration series (0%, 15%, 25%, 40%, 50%, 65% (v/v) of isotone NaCl solution in H_2_O and 0.9% NaCl solution in D_2_O) was measured. Areas below absorption bands were blotted against water concentration and linear regression was calculated through data points to obtain the slope (*m*) and *y*‐ interception (*y*_0_), to calculate water concentrations as described in detail before ^70^. The apical volume (*V*_api_) was calculated according to:

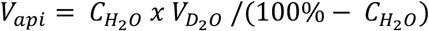

with 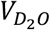 being the volume (25 *μ*l) of D_2_O in 0.9% NaCl. Changes in apical volume (Δ*V*_*api*_) were calculated by subtracting the remaining volume from the initial volume (25 µl isotonic NaCl solution). Changes in apical volumes in experimental filters were corrected for changes in evaporation controls.

### Infant mouse studies

7-day old C57BL/6 mice were used in accordance with the University of Maryland, Baltimore Institutional Animal Care and Use Committee. A streptomycin resistant derivative of the Tohama I “wild-type” *B. pertussis* strain was grown on Bordet-Gengou (BG) agar plates supplemented with defibrinated sheep blood (10%, Lampire Biological Products, Pipersville, Pennsylvania, USA) and 200 μg/ml streptomycin (Sigma-Aldrich, Merck). Animals were challenged via aerosol administered by a nebulizer system (Pari Vios) for twenty minutes. Lungs were excised post-euthanasia, homogenized mechanically (Omni International, Tulsa, Oklahoma, USA) and bacterial burden determined by plating on BG agar. Animals were intranasally administered vehicle (Cremophor:Ethanol:Saline 10:1:89), CsA (25 mg/kg, Sigma-Aldrich, Merck) or NIM811 (25 mg/kg, MedChemExpress, Monmouth, New Jersey, USA). To measure leukocytosis blood was harvested 7 days post-infection by cardiac puncture into ethylenediaminetetraacetic acid (EDTA) containing 1.7mL tubes (Fisher Scientific) and treated with ammonium-chloride-potassium (ACK) lysis buffer to lyse red blood cells. White blood cells were then counted using a hemocytometer.

### Ethics statement

Animals were used in accordance with the University of Maryland Institutional Animal Care and Use Committee protocol 0417005 (University of Maryland, Baltimore, MD) following approval by the Office of Animal Welfare Assurance. Animals were euthanized by CO_2_ asphyxiation followed by cervical dislocation and thoracotomy. All methods were performed in accordance with the relevant guidelines and regulations.

## Acknowledgements

This work was supported by the Deutsche Forschungsgemeinschaft (grant BA2087/2-2 to H.B.) and the Medical Faculty Ulm (Baustein 3.2 to K.E.). This project is supported by the European Social Fund and by the Ministry of Science, Research and the Arts Baden-Württemberg (fellowship to K.E.). V.W. is a fellow of the International Graduate School in Molecular Medicine Ulm (IGradU) and N.E. and R.L. were supported by the “Promotionsprogramm Experimentelle Medizin” of the Medical Faculty Ulm. Cordelia Schiene-Fischer is thanked for providing Cyp, FKBP and Hsp/c70 proteins, Johannes Buchner for providing Hsp90 protein, and Felix Hausch for FKBP FK1 fragments.

## Author contributions

KE designed the study, supervised and conducted experiments and wrote the manuscript. AKM, VW, NE, AA, MS, RL and JW conducted experiments. CS and NHC designed and conducted experiments and wrote parts of the manuscript. MF designed experiments and wrote parts of the manuscript. HB designed and supervised the study and worked on the manuscript.

