## Supplemental Figures 1+2 for "Pharmacological targeting of host chaperones protects from pertussis toxin in vitro and in vivo"

Running title: Chaperones facilitate pertussis toxin uptake into cells

*Katharina Ernst: Institute of Pharmacology and Toxicology, University of Ulm Medical Center, Ulm, Germany;, Tel. +49 731 50065528, Fax. +49 0731 50065502; Holger Barth: Institute of Pharmacology and Toxicology, University of Ulm Medical Center, Ulm, Germany;, Tel. +49 731 50065503, Fax. +49 0731 50065502

[#]These authors contributed equally to this work

**Keywords:** bacterial toxins, intracellular transport, chaperones, pharmacological inhibitors, human primary epithelium, pertussis toxin, Bordetella

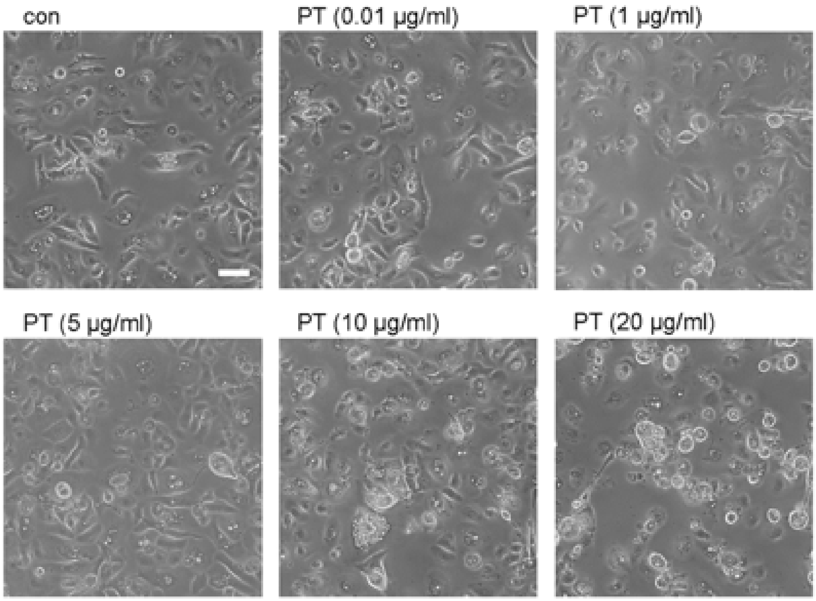


**S1 Figure. Effect of PT on human primary basal cells from bronchial epithelium.** Basal cells were incubated with different concentrations of PT or left untreated for control. Images were obtained after 25 h of PT treatment. Scale bar = 50 µm.

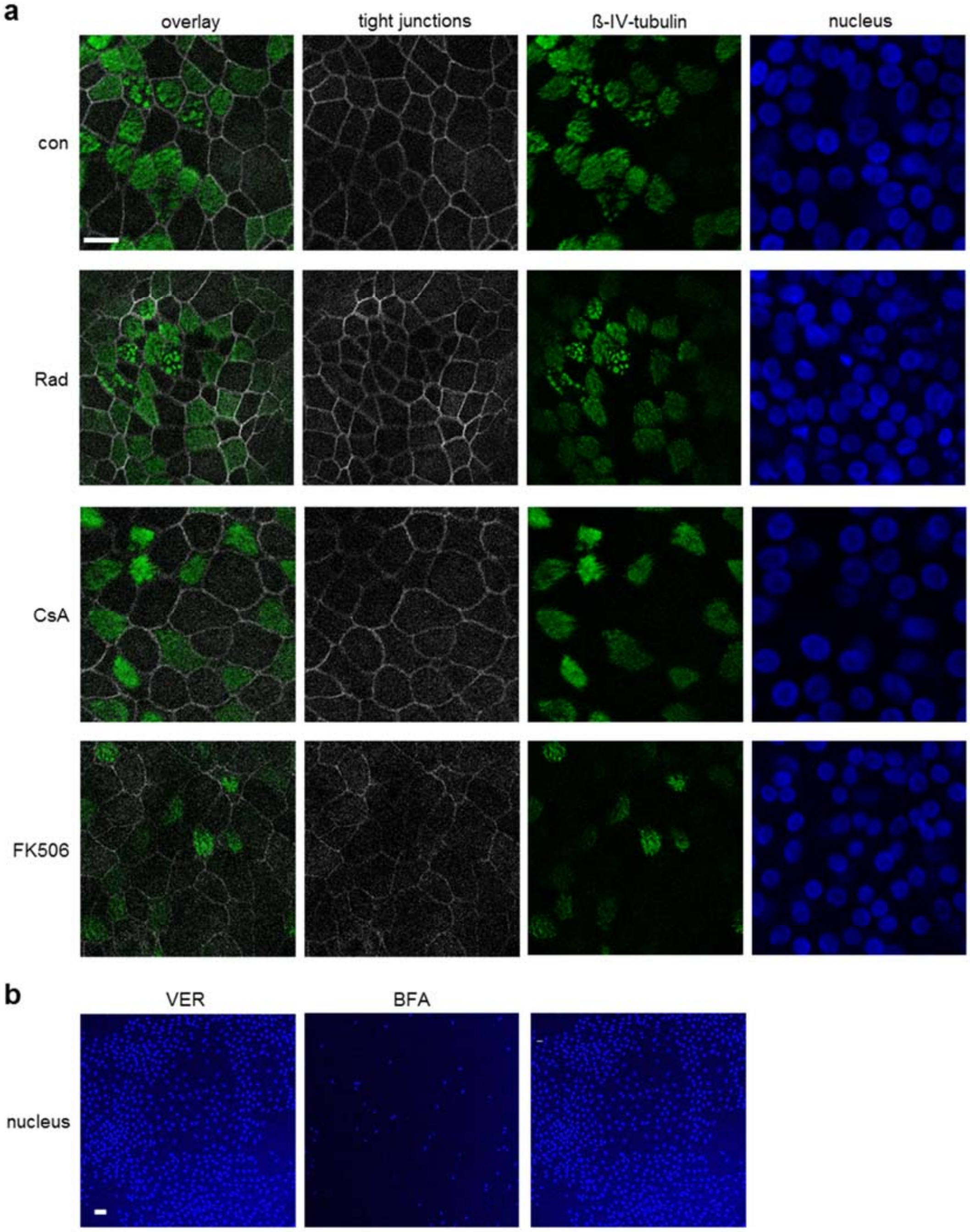


**S2 Figure. Effect of chaperone inhibitors on tight junctions of hBAECs.** hBAECs were incubated with Rad (20 µM), CsA (20 µM), FK506 (20 µM) **(A)**, VER (30 µM) and BfA (20 µM) **(B)** from basolateral side for 72 h or left untreated for control. Cells were fixed with 4 % PFA. For permeabilization and quenching of autofluorescence, cells were treated with 0.2 % saponin. The nuclei were stained with Hoechst33342 (blue) and the tight junctions were stained with ZO-1 (grey). Type IV ß-tubulin (green) was stained with a specific primary antibody and the respective fluorescence-labeled secondary antibody. Pictures were taken with an inverted confocal microscope. Scale bar = 20 µm **(A)**, 40 µm **(B)**.
